# Elucidating host cell response pathways and repurposing therapeutics for SARS-CoV-2 and other coronaviruses using gene expression profiles of chemical and genetic perturbations

**DOI:** 10.1101/2022.04.18.488682

**Authors:** Zhewei Shen, Anna Halberg, Jia Yi Fong, Jingyu Guo, Gavin Song, Brent Louie, Gregory R. Luedtke, Viwat Visuthikraisee, Andy Protter, Xiaoying Koh, Taegon Baik, Pek Yee Lum

**Author notes:** Contributed equally (Zhewei Shen, Anna Halberg, Jia Yi Fong, Jingyu Guo).

## Abstract

COVID-19 is an ongoing pandemic that has been causing devastation across the globe for over 2 years. Although there are multiple vaccines that can prevent severe symptoms, effective COVID-19 therapeutics are still of importance. Using our proprietary *in silico* SMarTR™ engine, we screened more than 22,000 unique compounds represented by over half a million gene expression profiles to uncover compounds that can be repurposed for SARS-CoV-2 and other coronaviruses in a timely and cost-efficient manner. We then tested 13 compounds *in vitro* and found three with potency against SARS-CoV-2 with reasonable cytotoxicity. Bortezomib and homoharringtonine are some of the most promising hits with IC_50_ of 1.39 μM and 0.16 μM, respectively for SARS-CoV-2. Tanespimycin and homoharringtonine were effective against the common cold coronaviruses. In-depth analysis highlighted proteasome, ribosome, and heat shock pathways as key targets in modulating host responses during viral infection. Further studies of these pathways and compounds have provided novel and impactful insights into SARS-CoV-2 biology and host responses that could be further leveraged for COVID-19 therapeutics development.

## Introduction

In December 2019, a new coronavirus emerged and rapidly spread around the world, causing a global pandemic on an unprecedented scale. SARS-CoV-2, the virus that causes COVID-19, is a positive-sense single-stranded RNA coronavirus.

Due to the virus’s rapid transmission rate, asymptomatic infection, and the emergence of multiple infectious variants, the timely development of effective therapeutic modalities has been a global challenge. While efforts in vaccine development have resulted in emergency use authorization of multiple effective vaccines and the recent approvals of coronavirus targeting anti-viral compounds, we believe that understanding the biology of infection and the host cell responses will allow us to develop novel therapeutics that could be used concomitantly with current antivirals and vaccines to prevent severe diseases. In addition, we also studied other common cold coronaviruses towards the goal of developing pan-coronavirus therapeutics that could slow or stop viral production in host cells.

We used gene expression profiles of both compound and genetic perturbations as probes for understanding the biology of viral replication. Using this approach, we predicted for compounds that may impede viral production in the host cells. Given SARS-CoV-2 is closely related to SARS-CoV-1 and are both from the beta coronavirus family [1,2] and the abundance of SARS-CoV-1 data, we used gene expression from viral studies of both viruses for our predictions. The approach starts with our proprietary computational engine (SMarTR™ engine) that analyzes gene expression data of disease cells and chemical and genetic perturbations to generate *in silico* predictions for possible treatments.

Towards this goal, we screened *in silico* more than 22,000 existing compounds represented by over half a million gene expression profiles, rapidly producing candidate compounds that can be further tested. In summary, we found that chemically perturbing translation initiation or regulation, heat shock, and proteasomal pathways, were effective in controlling viral reproduction in a pan-virus or virus-specific manner. Small molecules like bortezomib, homoharringtonine and tanespimycin that affected these pathways were able to interfere with viral production against SARS-CoV-2 or the common cold viruses HCoV-229E and HCoV-OC43.

## Results

### Gene expression based *in silico* “transcriptotypic screen” approach

Because of the genomic similarity between SARS-CoV-1 and SARS-CoV-2, we used gene expression profiles of human lung cell lines and peripheral blood from patients infected with either SARS-CoV-1 or SARS-CoV-2 for the predictions. Host cell response profiles from each condition were used independently for the prediction and tested against a library of gene expression profiles of cell responses to over 22,000 unique compounds or gene perturbations individually.

From the initial raw predictions, we executed our first-round selection by ranking the compounds based on the number of times a compound passes the Fisher exact test in each input data independently. We believe this stringent criteria allows us to identify compounds with the strongest potential to affect viral production. We identified a total of 1416 such ranked compounds from over 6,700 potential compounds that can be sourced and therefore tested quickly. In this round of testing, priority was given to groups of compounds representing the same classes of mechanisms of action (MOA) if multiple of which are top ranked. We then added an additional filter to exclusively focus on launched or clinical phase compounds and tested a representative candidate from each MOA of interest. A total of five compounds, namely trametinib (MEK inhibitor), bortezomib (proteasomal inhibitor), homoharringtonine (protein synthesis inhibitor), dasatinib (Bcr-Abl kinase inhibitor), and lacidipine (calcium channel blocker) were selected for testing in SARS-CoV-2 infected Vero cells. Selective index (SI) was used as a measurement of a compound’s therapeutic benefit over risk.

Of the aforementioned compounds, trametinib (SI 3.34) and bortezomib (SI 35.9 and SI 14.63) showed anti-SARS-CoV-2 activities and cytotoxicity over 50 μM (Table 1, Figure 1a and 1b). Homoharringtonine is highly effective against SARS-CoV-2 but also exhibits some level of toxicity. However, the SIs of homoharringtonine (12.84 and 24.06, Table 1, Figure 1a and 1b) suggest that there may be a reasonable therapeutic window for anti-SARS-CoV-2 activity before cytotoxicity is induced.

**Figure 1:**
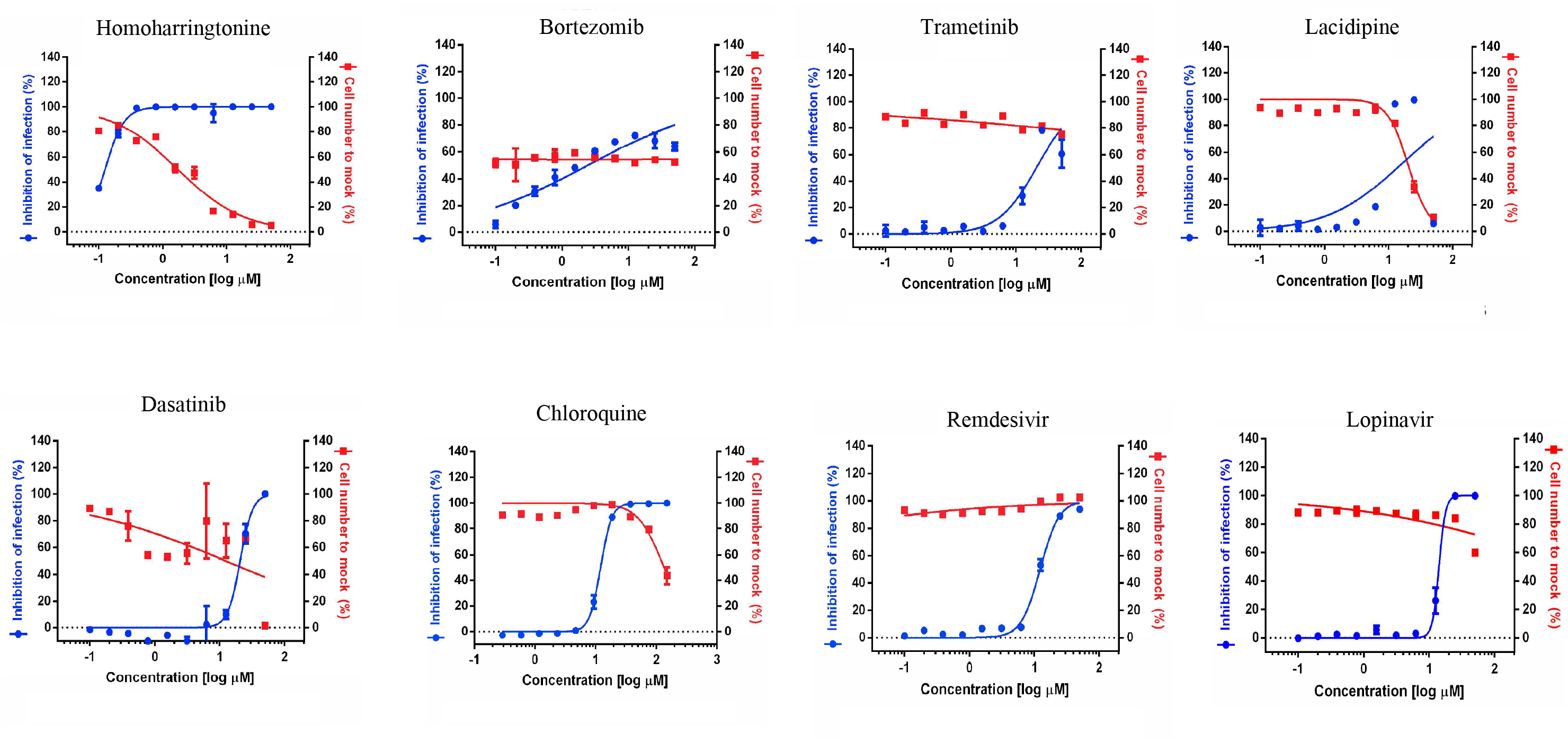
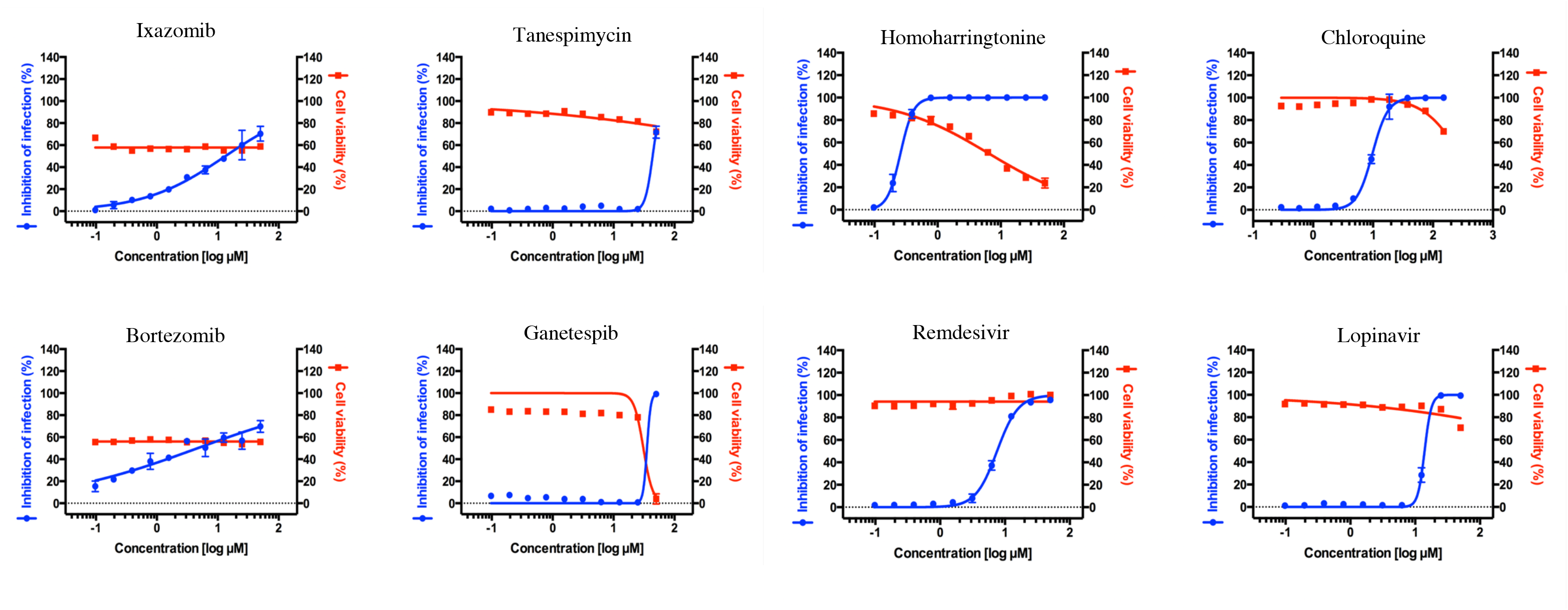
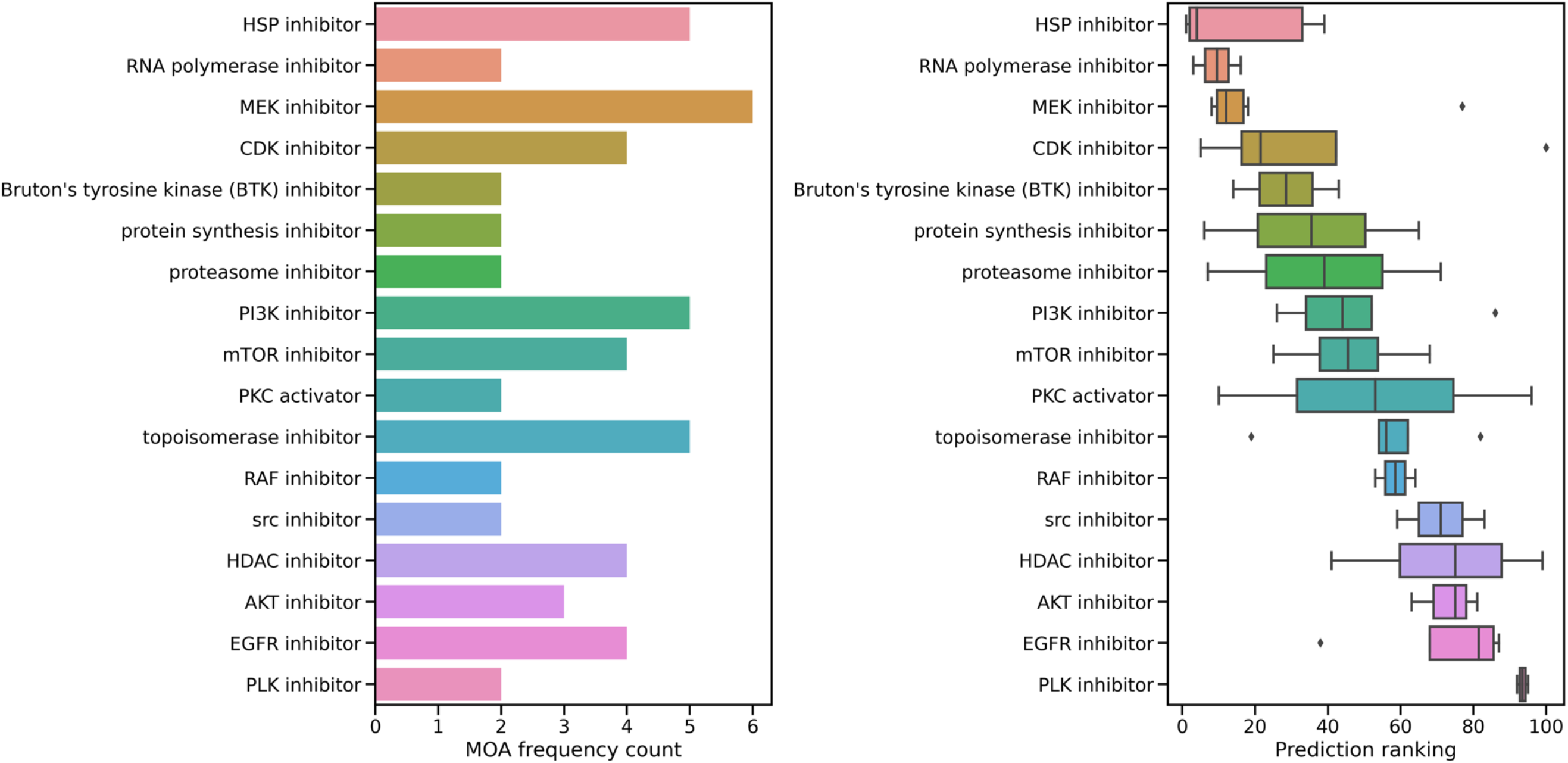
(A) Viral inhibition and cell viability curves used to generate Table 1 Round 1. (B) Viral inhibition and cell viability curves used to generate Table 1 Round 2. (C) Comparison of drug MOA class frequency count and predicted compound prediction ranking in Top 100 compounds with strongest predicted anti-viral strength.

**Table 1:** Predicted anti-SARS-CoV-2 compounds *in vitro* efficacy testing from two independent rounds of testing. IC_50_, CC_50_, and SI index were calculated of each compound in SARS-CoV-2 infected in Vero cells.

|  | **Compound** | **IC<sub>50</sub> (μM)** | **CC<sub>50</sub> (μM)** | **SI index** |
| --- | --- | --- | --- | --- |
| Round 1 | Homoharringtonine | 0.16 | 2.14 | 12.84 |
|  | Bortezomib | 1.39 | >50 | 35.93 |
|  | Trametinib | 14.95 | >50 | 3.34 |
|  | Lacidipine | 14.89 | 20.15 | 1.35 |
|  | Dasatinib | 19.74 | 13.20 | 0.67 |
|  | Remdesivir | 12.08 | >50 | 4.14 |
|  | Chloroquine | 12.0 | 127.8 | 10.65 |
| Round 2 | Ganetespib | 35.50 | 28.95 | 0.82 |
|  | Ixazomib | 11.71 | >50 | 4.27 |
|  | Tanespimycin | 31.33 | >50 | 1.60 |
|  | Bortezomib | 3.42 | >50 | 14.63 |
|  | Homoharringtonine | 0.26 | 6.22 | 24.06 |
|  | Chloroquine | 9.80 | >150 | 15.31 |
|  | Remdesivir | 7.45 | >50 | 6.71 |
|  | Lopinavir | 13.50 | >50 | 3.70 |

The next set of compounds tested was comprised of top ranked compounds that perturb the same pathways. We tested two strongly predicted proteasomal inhibitors namely bortezomib, that showed anti-SARS-CoV-2 activity in the previous experiment, and its analog ixazomib. We also selected two highly ranked compounds from the heat shock pathway, namely tanespimycin and ganetespib. Both the proteasomal pathway and the heat shock pathway were significantly represented among the top 100 ranked compounds (Figure 1c). Consistent with the initial study, bortezomib and ixazomib showed anti-SARS-CoV-2 activity with a SI of 14.63 and 4.27 respectively (Table 1). Tanespimycin and ganetespib, on the other hand, showed no activity against SARS-CoV-2 in a Vero cell assay despite having very strong prediction scores from the engine (Table 1, Figure 1b). The potent anti-viral activity of homoharringtonine was consistent across the two independent studies (Figure 1a and 1b).

As another test elucidating the biology of host responses to SARS-CoV-2, we identified several compounds that were weakly predicted (poorer ranks) but with novel MOAs. Meclofenamic acid (nonsteroidal anti-inflammatory), sitagliptin (DPP4 antagonist), and levetiracetam (anticonvulsant, MOA unclear) did not show meaningful anti-SARS-CoV-2 activity while SR-2640 (leukotriene D4 and E4 receptor antagonist) and AZD-1208 (PIM kinase inhibitor) had very poor SI values (Table 2, Figure 2).

**Figure 2:**
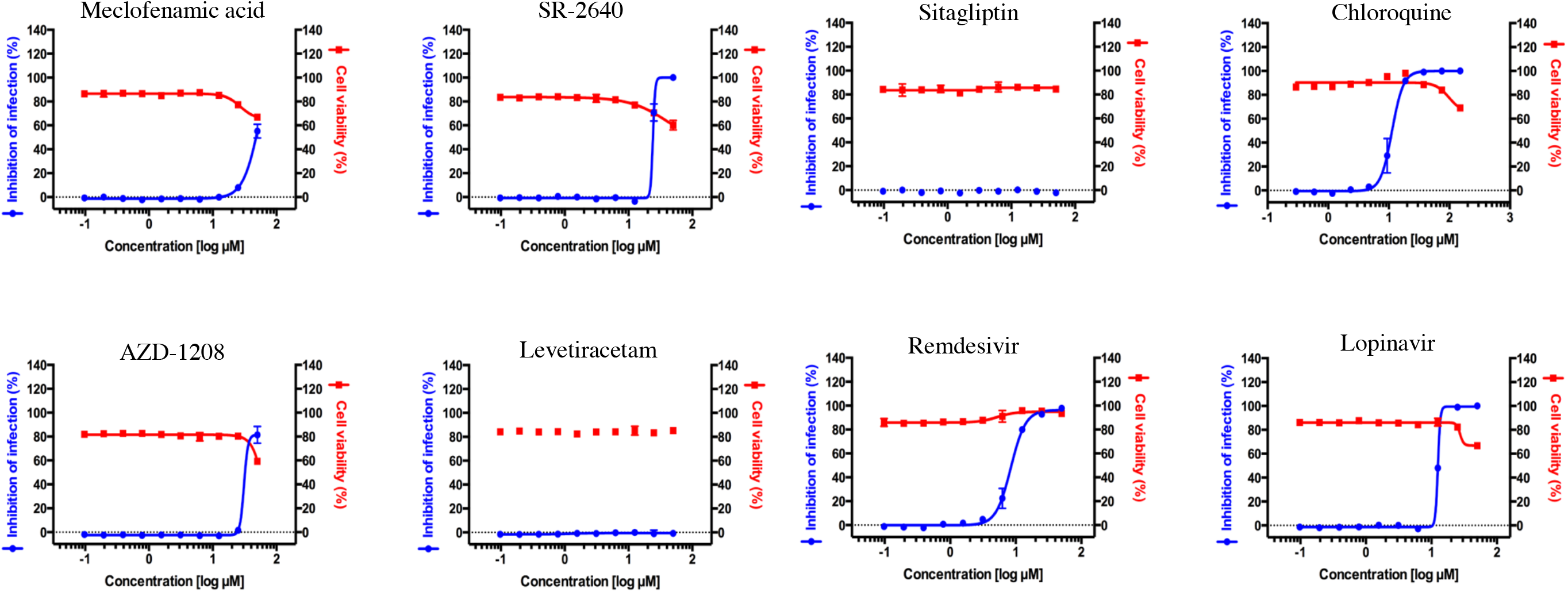
Viral inhibition and cell viability curves used to generate Table 2.

**Table 2:** Average reversal score predicted anti-SARS-CoV-2 compounds *in vitro* efficacy testing. IC_50_, CC_50_, and SI index were calculated of each compound in SARS-CoV-2 infected in Vero cells.

| **Compound** | **IC<sub>50</sub> (μM)** | **CC<sub>50</sub> (μM)** | **SI index** |
| --- | --- | --- | --- |
| AZD-1208 | 27.34 | >50 | 1.83 |
| Sitagliptin | >50 | >50 | 1.00 |
| SR-2640 hydrochloride | 24.23 | >50 | 2.06 |
| Levetiracetam | >50 | >50 | 1.00 |
| Meclofenamic acid sodium salt | 35.13 | >50 | 1.42 |
| Chloroquine | 11.29 | >150 | 13.29 |
| Lopinavir | 12.63 | >50 | 3.96 |
| Remdesivir | 8.58 | >50 | 5.83 |

### Efficacy of compounds against common cold-causing human coronaviruses

We were also interested to understand if these compounds and their target pathways have wider applications in modulating viral host responses, specifically other coronaviruses. The two common cold viruses, HCoV-229E and HCoV-OC43, belong to the coronavirus alpha and beta subgroup, respectively. Both SARS-CoV-1 and SARS-CoV-2 are also in the beta subgroup [1,2]. We tested bortezomib, tanespimycin, and homoharringtonine against these two cold viruses in a human lung fibroblast cell line. Our results show strain specific sensitivity to these compounds. Bortezomib showed no activity against HCoV-229E or HCoV-OC43 in three independent studies (Table 3, Figure 3), though it has favorable SIs against SARS-CoV-2 (Table 1). Tanespimycin, on the other hand, was highly effective against HCoV-229E and HCoV-OC43 with an EC_50_ of less than 1 nM (Table 3, Figure 3), though it did not exhibit anti-SARS-CoV-2 activity in Vero cell studies (Table 1). Homoharringtonine was effective against HCoV-229E and HCoV-OC43 with EC_50_s < 100 nM (Table 3, Figure 3), as well as against SARS-CoV-2 with an average EC_50_/IC_50_ under 300 nM (Table 1).

**Figure 3:**
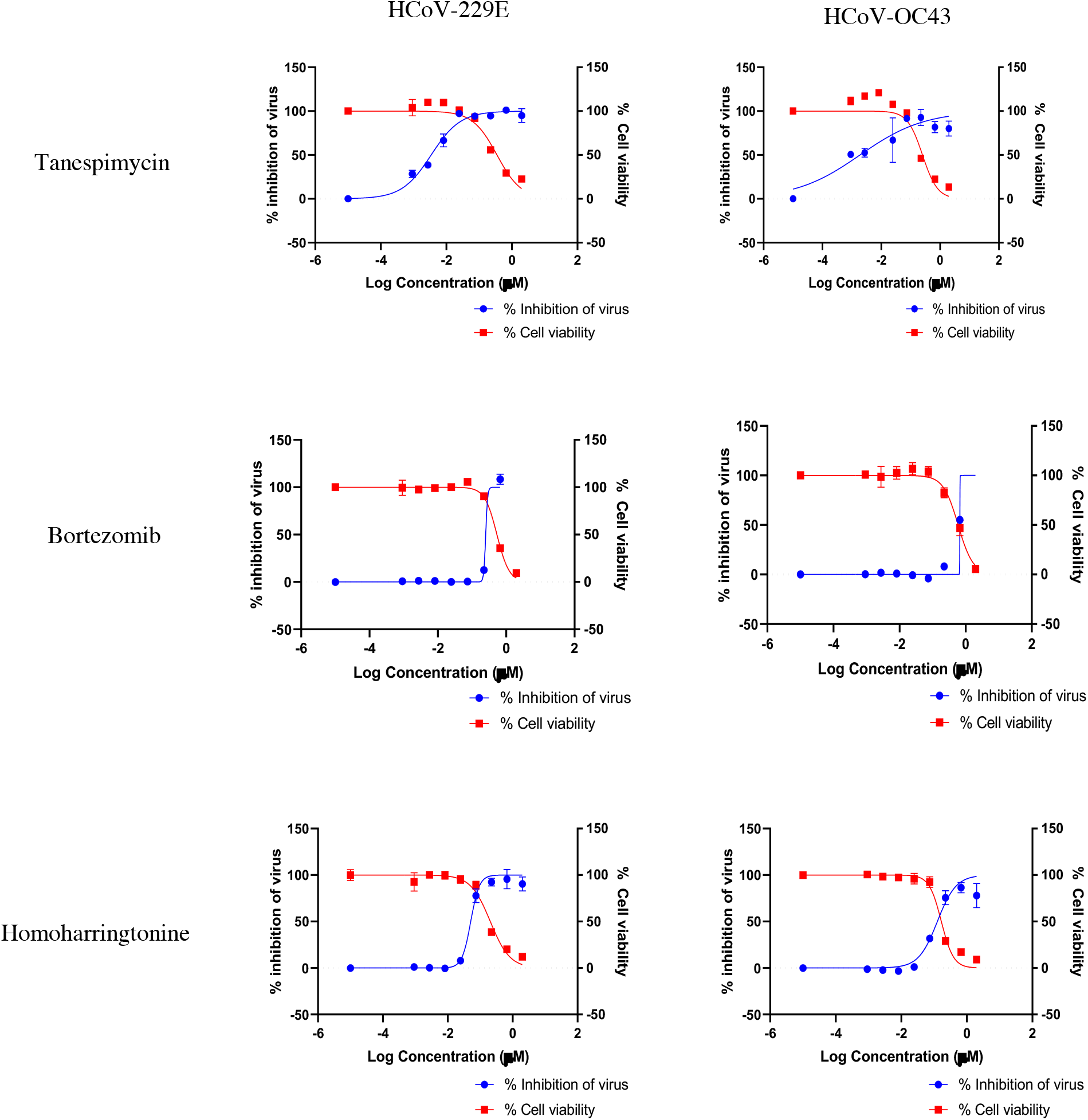
EC_50_ vs. CC_50_ curves associated with Table 3 for the 3 compounds tested against HCoV-229E and HCoV-OC43 in MRC5 cells

**Table 3:** (A) EC_50_, CC_50_, and SI index of 3 compounds of interest in MRC5 cells infected with HCoV-229E or HCoV-OC43.

| **HCoV strain** | **Compound** | **EC<sub>50</sub> (μM)** | **CC<sub>50</sub> (μM)** | **SI Index** |
| --- | --- | --- | --- | --- |
| HCoV-229E | Tanespimycin | 0.0036 | 0.182 | 101 |
|  | Bortezomib | 0.2518 | 0.5364 | 2 |
|  | Homoharringtonine | 0.05106 | 0.2046 | 4 |
| HCoV-OC43 | Tanespimycin | 0.0019 | 0.2528 | 133 |
|  | Bortezomib | 0.666 | 0.5914 | < 1 |
|  | Homoharringtonine | 0.1261 | 0.1670 | 1 |

### Pathways predicted that are amenable to small molecule intervention

In addition to the compounds we tested, we also investigated the pathways that are represented by the top 100 ranked predicted compounds. Of the top compounds with known MOAs, we found that compounds representing the heat shock, proteasomal, and protein synthesis pathways are ranked very highly in our approach in addition to being effective in curtailing viral production in our experiments (Figure 1c). In contrast, compounds representing the neurotransmitter related classes (e.g., serotonin receptor antagonists, dopamine receptor antagonists) and inflammatory (lipoxygenase, histamine, chemokine) are often represented by more lowly ranked compounds (Figure 4). As mentioned earlier, both levetiracetam and SR-2640 hydrochloride did not result in a meaningful effect on viral production (Table 2).

**Figure 4:**
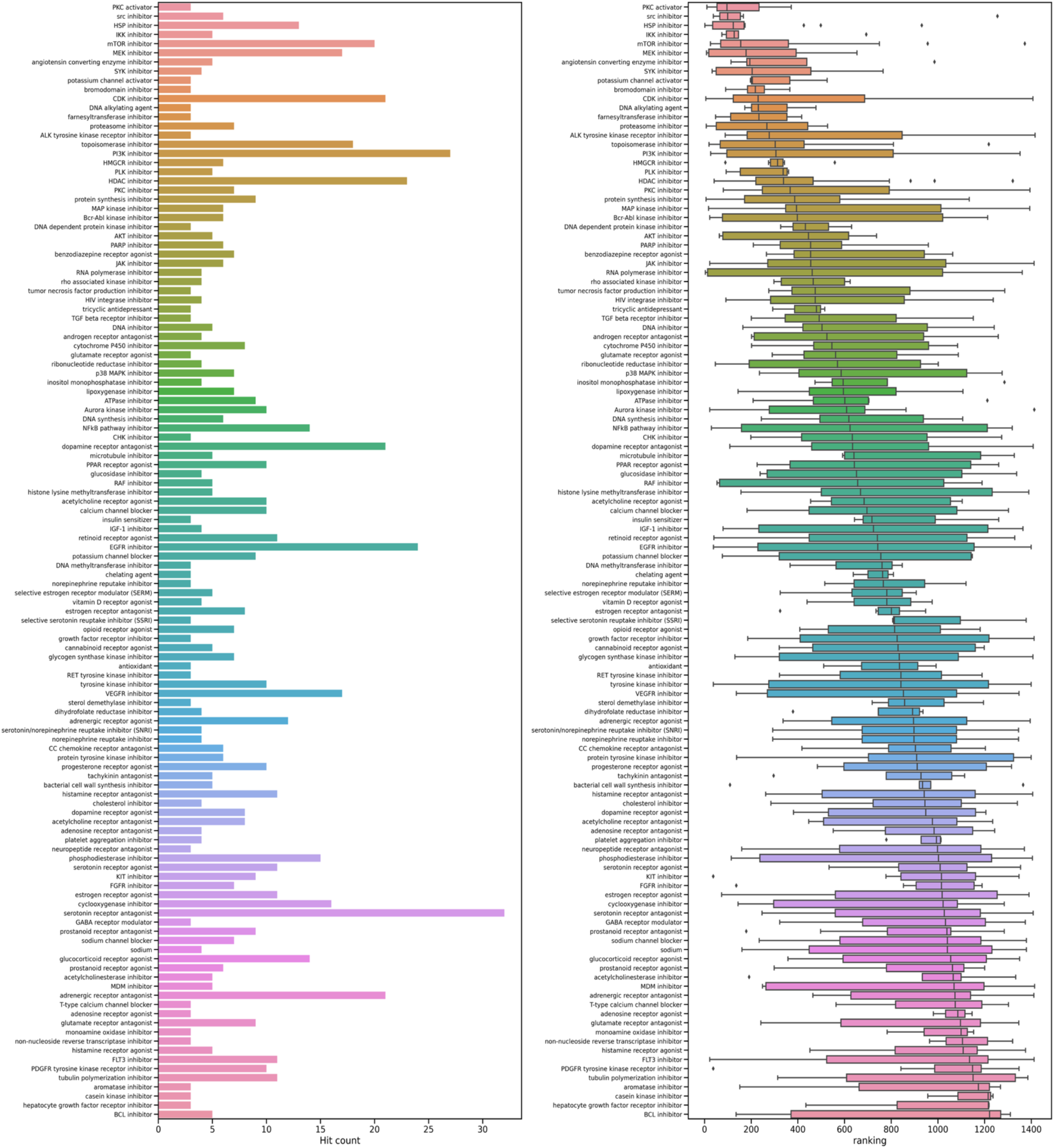
Visualization of drug class frequency count and prediction strength ranking of all predicted anti-SARS-CoV-2 compounds.

### Using expression profiles of gene knockdowns to probe host response biology

By processing the gene expression profiles of gene knockdowns in the same way as we did for compound-generated gene expression profiles, we generated a list of genes whose knocked-down gene expression profiles were predicted by the Auransa SMartTR™ engine to show strong anti-SARS-CoV-2 activities. Pathway analysis using predicted gene targets found a significant enrichment of proteasome pathway and heat shock pathway with p-values of 0.01 and 0.07 from hypergeometric tests, respectively. In fact, shRNA knockdown of HSP90AB1 is ranked the 14^th^ most likely candidate to reverse SARS-CoV-2 signatures in host cells.

### Deconvoluting the biology of the prediction

In order to better understand the MOA underlying the compounds that were predicted and showed efficacy, we explored the biological rationale underlying the prediction that yielded the three viable compounds, focusing on pathways that were deemed important by our engine as it made those predictions (Table 1). Our algorithm found that NF-kappaB signaling, proteasomal/ER stress activities, and cell cycle regulation are key targets affected by bortezomib. Tanespimycin shows *in silico* effects on heat shock pathways, oxidative stress responses, lipid metabolism, and cell cycle. Homoharringtonine has impacts on a bigger set of biological processes including cholesterol and steroid biosynthesis, cell cycle, and NF-kappaB signaling (Table 4).

**Table 4:** Selective pathways identified using top genes predicted by Auransa SMarTR engine™ to be reversed by anti-viral compounds based on STRING DB analysis.

| Compounds | Pathway name | Source | Strength | FDR |
| --- | --- | --- | --- | --- |
| Bortezomib | Canonical NF-kB pathways, and NF-kappa-B/Dorsal | STRING | 1.81 | 0.00055 |
|  | G2/M DNA replication checkpoint | REACTOME | 1.99 | 0.0267 |
|  | I-kappaB/NF-kappaB complex | GO-CC | 1.99 | 0.0018 |
| Homoharringtonine | I-kappaB/NF-kappaB complex | GO-CC | 1.92 | 0.0013 |
|  | Canonical NF-kB pathways, and NF-kappa-B/Dorsal | STRING | 1.81 | 0.00082 |
|  | Negative regulation of reactive oxygen species metabolic process | GO-BP | 1.23 | 0.0105 |
| Tanespimycin | Chaperone-mediated protein transport | GO-BP | 1.73 | 0.0071 |
|  | Cholesterol biosynthesis | UniProt | 1.47 | 0.0137 |
|  | Steroid biosynthesis | UniProt | 1.65 | 0.00114 |

## Discussion

We used our proprietary computational engine to predict compounds and targets in order to study host cell responses to infections by SARS-CoV-1, SARS-CoV-2 and 2 common cold coronaviruses. We showed that the most strongly predicted compounds belong to the classes of compounds known to modulate the following pathways: PKC, SRC, HSP, IKK, mTOR, ACE and proteasome (Figure 4). Interestingly, the most frequently predicted drug class among the 1416 unique compounds are the neurotransmitter related classes (e.g., serotonin receptor antagonists, dopamine receptor antagonists). However, these compounds generally have lower rankings compared to compounds from the heat shock, proteasomal and gene translation classes (Figure 4). Levetiracetam, one of the compounds in the most frequent but low-ranking category, had no anti-viral activity (Table 2, Figure 2). In contrast, the positive results we obtained are from highly ranked predicted compounds belonging to the cell growth, proteasome, and heat shock drug classes. These results suggest that frequency of drug classes may not be as important for compound selection as much as the predicted ranks that are based on the engine’s disease reversal strength scores (Figure 4). Counting drug class frequency also has the bias of more frequently annotated, perhaps older and larger classes of popular compounds compared to newer or smaller classes of compounds.

Many compounds have been predicted or tested for potential activities against SARS-CoV-2 since the start of the pandemic by other research groups. Numerous early- and late-stage ongoing clinical trials are focused on compounds mostly belonging to the anti-inflammation or direct anti-viral growth classes [3]. Our engine identified some of these reported anti-SARS-CoV-2 compounds, specifically emetine (protein synthesis inhibitor), niclosamide (anthelminthic), and pralatrexate (antineoplastic folate analog metabolic inhibitor) [4–6]. Niclosamide and emetine showed impressive anti-SARS-CoV-2 capabilities in published preclinical studies, with reported SI index of 178.57 and 176.65, respectively. They are ranked #6 and #28 in our predictions. Pralatrexate, with a reported IC_50_ of 0.008 μM [6], is ranked within the top 60%.

One interesting pathway in our *in silico* prediction is the proteasome pathway. The proteasome inhibitors bortezomib and ixazomib showed efficacy in controlling viral load without significant cytotoxicity. Of the two, bortezomib exhibited more potency. The ubiquitin-proteasome pathway is a key pathway regulating a variety of cellular processes. Viruses are known to hijack this pathway for their propagation [7]. Interestingly, we did not find bortezomib to be effective against the cold viruses. It is unclear as to why ubiquitin-proteasome pathway inhibition had no effect in our anti-cold viral assays. One reason could be virus specific susceptibility to proteasome inhibition. We also cannot rule out cell type specificity having an impact on the results of the *in vitro* assay. Furthermore, whether proteasome inhibitors can be developed as an anti-viral depends on their efficacy vs. toxicity profiles *in vivo*. We took up the exercise of calculating the clinical exposure of bortezomib from publicly known clinical data and found it to be approximately 580 nM via intravenous infusion for cancer (see Materials and Methods). IC_50_s of bortezomib in the *in vitro* assays are 1.39 μM and 3.42 μM with CC_50_ > 50 μM, indicating that there may be a reasonable window of safety.

Several publications have also mentioned bortezomib as a potential candidate either by different *in silico* approaches or via compound screening in the laboratory. The *in silico* approaches ranged from prediction only to analyzing transcription factors regulatory and protein-protein interactions networks. Pan *et al.* and Adhami *et al.* used preselected gene sets representing viral responses to predict anti-COVID-19 compounds [8,9]. Xing *et al.* utilized known antivirals as internal comparisons to identify and test compounds that can significantly reverse COVID-19 viral signature [10]. These different approaches all led to bortezomib as a candidate for anti-SARS-CoV-2. These studies, together with our unsupervised (without a known training set, preselection of genes, or previously tested positive compound ‘hits’) *in silico* approach, further validate the proteasomal pathway as a critical pathway that can be targeted as an anti-viral strategy.

The next pathway of significance in our prediction is RNA translation. We predicted and validated *in vitro* that ribosome inhibitor homoharringtonine is highly effective against both SARS-CoV-2 and cold coronaviruses. Schubert *et al.* showed that the Nsp1 (also known as the host shutoff factor) interferes with the ribosomal mRNA channel to stop translation of host mRNAs as a host defense against SARS-CoV-2 infection [11]. Although targeting something as critical as RNA translation may adversely impact the host cells, we believe that homoharringtonine, an inhibitor of the ribosome complex, should be further studied as a potential antiviral drug. Even though stopping RNA translation can be detrimental for both the host cell and virus, SIs of homoharringtonine (12.84 and 24.06, Table 1, Figure 1a and 1b) indicate that there might be enough of a window to give the host cell an edge by stopping the very mechanisms that the virus relies on to replicate. Other reports using *in vitro* cell based screening of FDA approved drugs such for TMPRSS2 reduction or *in silico* prediction to affect IFN-beta genes also identified homoharringtonine as a promising candidate [12,13]. Similar to bortezomib, these studies, together with the independent identification by our engine, indicates that gene translation is a critical machinery used by the virus to replicate. We also show that homoharringtonine is potent against HCoV-229E and HCoV-OC43 with EC_50_s < 100 nM, indicating perhaps RNA translation may be a common theme that can be exploited as a pan anticoronavirus strategy. In addition, we calculated the clinical exposure of homoharringtonine to approximately 66 nM. IC_50_s of homoharringtonine is 0.16 μM and 0.26 μM in our testing (see Materials and Methods), hence we also believe that additional effort is warranted to explore homoharringtonine as a potential pan anti-coronavirus compound. It is also worth adding that a clinical trial in COVID-19 patients treated with nebulized homoharringtonine is ongoing [14].

The last compound, tanespimycin, was predicted very strongly from the SARS-CoV-1 and SARS-CoV-2 data but surprisingly, neither tanespimycin nor a second generation HSP90 inhibitor ganetespib exhibited anti-viral efficacy in our SARS-CoV-2 assays. In contrast, tanespimycin was highly potent against the 2 cold coronaviruses. Interestingly, Li *et al.* have shown that 10 μM tanespimycin effectively inhibited viral activities of SARS-CoV-1, SARS-CoV-2, and MERS-CoV in Huh-7 cells [15]. We believe that the HSP90 pathway deserves a closer look in other cell types and conditions. Furthermore, we postulate that discrepancy in tanespimycin *in vitro* efficacy against SARS-CoV-2 may be partly caused by the difference in cell models used (i.e., Vero vs. Huh-7). Prior publication has eluded to anti-SARS-CoV-2 drug response disparity in a cell line dependent manner in Vero and Calu-3 cells [16].

In 2020, remdesivir, a viral RNA polymerase inhibitor originally developed to treat Ebola patients, was found to show clinical improvement in COVID-19 patients in clinical trials and subsequently approved in the United States in October 2020 [17]. We note that remdesivir was not predicted by our approach. This is likely because our approach uses gene expression profiles of infected host cells to predict compounds rather than targeting the viral proteins. Remdesivir, on the other hand, targets the viral RdRp, with many fold specificity over human RNA Polymerase II and mitochondrial RNA polymerase [18–20], hence, we do not expect to see remdesivir or any other virus-targeting compounds to be predicted using our approach.

In summary, our approach has pointed us to several critical host cell pathways that could be targeted to stop coronavirus replication, namely the proteasomal pathway, protein synthesis or post-translational regulation machinery, and the heat shock system. We identified compounds in the above pathways that are candidates for repurposing. Although the safety profiles of these potential candidates need to be considered in an anti-viral context, we believe that repurposing drugs with clinical approval or in advanced clinical phases with acceptable clinical safety will greatly shorten the development time and provide the opportune therapies during an ongoing pandemic. In fact, several existing drugs had been recommended to treat hospitalized COVID-19 patients, including antiviral drugs (e.g., remdesivir), anti-inflammatory drugs (e.g., baricitinib and corticosteroids), and intravenous monoclonal antibodies [21].

We demonstrated that an *in silico* approach such as ours, done without pre-defined sets of genes or the necessity to train using existing known anti-viral agents, can generate promising anti-viral compounds and vulnerable pathway predictions, with *in vitro* efficacy and reasonable cytotoxicity. Importantly, our prediction focuses us on compounds and pathways that can modulate host responses in a more pan-coronavirus way instead of inhibiting specific viral proteins. These compounds may have wider clinical applications beyond SARS-CoV-2 treatment as shown in our *in vitro* results and are less constrained by the virus strains as exemplified by homoharringtonine. We believe that our work shows strong support for antiviral therapy development focused on host response regulation.

## Materials and Methods

### Data curation and processing

SARS-CoV-1 and SARS-CoV-2 studies used in the analysis were curated from NCBI Gene Expression Omnibus (GEO). The following datasets were used: GSE17400, GSE47960, GSE47961, GSE47962, GSE1739, GSE5972, PRJNA625518, PRJNA631969, and PRJNA637580.

Microarray data were directly downloaded from GEO and log2 transformed where necessary. RNAseq data were processed using Salmon algorithm which was adapted to Auransa’s internal pipelines [22]. Log fold change (Log2FC) values were computed by contrasting a total of 14 selected viral infection conditions against corresponding controls.

Auransa’s curated gene expression database of compounds and genetic perturbations comprise of over half a million gene expression profiles across over 22K unique compounds (drug induced gene expression signature, DIGS). The DIGS data were used to assess compound reversal strength against the gene expression profiles generated from human cells infected with viruses of interest. All publicly available gene expression profiles were downloaded from the GEO.

### Computational compound prediction

For compound prediction, we designed our prediction algorithm to score and select the compounds that maximize the reversal effect of the human genomic expression under virus infection, such that these compounds may have the potential to correct the patient phenotypes under virus infection. This algorithm is, in part, based on the concept of GSEA [23]. All genes in the gene expression profiles were used in the algorithm, without any pre-selection that might bias the results. Our algorithm also does not need to be trained on any pre-existing list of known antiviral compounds, hence opening us up to discovering compounds and critical pathways in an unsupervised manner.

After the reversal scores were calculated, we used a Fisher Exact test-based method for ranking. The Fisher Exact test was used to examine if any single compound has a significant good-hit-bad-hit count ratio compared to all other compounds within the same DIGS. FDR correction was performed and filtered at the significance threshold of 0.05. Each time a compound has a significant FDR value is considered a hit count of 1. Finally, the total number of counts across all input conditions is summed into a single cumulative sum for ranking. To break the ties in ranking, we also calculate the percentage of number of drug signatures fulfilling the reversal score threshold / total number of drug signatures for a single compound for contrasts where the Fisher Exact test yielded a significant result.

We also tested compounds that may not pass the method above but nonetheless exhibited on average a good reversal score across 14 contrasts on the drug signature level and is of novel MOAs. An average reversal score was computed for each drug signature and then filtered using the predetermined reversal score threshold. These drug signature conditions were ranked based on the average score in an ascending manner. We selected a few compounds in this category to represent each pathway of interest.

### Pathway enrichment analysis

Top 100 most significant genes identified by Auransa SMarTR engine™ to be reversed by predicted anti-viral treatment were analyzed using STRING database [24,25] (STRING DB, https://string-db.org/). The genes used for pathway analysis are listed in Supplementary Table 4.

### Compound sourcing

All compounds are research grade chemicals sourced from MedChemExpress, Sigma-Aldrich, Tocris Bioscience, and Selleck Chemicals.

### Anti-SARS-CoV-2 compound testing

Predicted compounds were tested by the Institut Pasteur Korea (Seongnam, South Korea). SARS-CoV-2 virus was used to infect Vero cells at an MOI of 0.0125 for 24 hours after a one-hour preincubation with the compounds. Cells were fixed with 4% paraformaldehyde (PFA), followed by permeabilization. Subsequently, cells were stained with anti-SARS-CoV-2 Nucleocapsid (N) primary antibody, a 488-conjugated goat anti-rabbit IgG secondary antibody and Hoechst 33342. Fluorescence expression was imaged using Operetta (Perkin Elmer), a large-capacity image analysis device. The acquired images were analyzed using an institutional proprietary Image Mining (IM) software. The total number of cells per well was calculated as the number of nuclei stained with Hoechst, and the number of infected cells was calculated as the number of cells expressing the viral N-protein. Infection ratio was calculated as the number of cells/total cells expressing the N protein. Dose response curve values were computed using the equation: Y =Bottom + (Top Bottom) / (1 + (IC50/X)Hillslope) using XLFit 4 (IDBS) software. All IC_50_ and CC_50_ values were measured in two repeated experiments, and the reliability of the assay was verified by the values of the Z-factor and the coefficient of variation (%CV).

Cytopathic effect of viral infection was measured using microscopy after 3 days of infection. Plates were stained with neutral red dye for approximately 2 hours (±15 minutes). Supernatant dye was removed, and wells rinsed with PBS. The incorporated dye was extracted in 50:50 Sorensen citrate buffer/ethanol for >30 minutes and the optical density was read on a spectrophotometer at 540 nm. Optical densities were converted to percent of cell controls and normalized to the virus control, then the concentration of test compound required to inhibit CPE by 50% (EC_50_) was calculated by regression analysis. The concentration of compounds that would cause 50% cell death in the absence of virus was similarly calculated (CC_50_). The selective index (SI) is the CC_50_ divided by EC_50_.

For virus yield reduction assays, the supernatant fluid from each compound concentration was collected on Day 3 post-infection, before neutral red staining and tested for virus titer using a standard endpoint dilution CCID50 assay and titer calculations using the Reed-Muench (1948) equation. The concentration of compounds required to reduce virus yield by 1 log10 (EC_90_) was calculated by regression analysis. A selectivity index (SI index) was calculated for each compound as SI index = CC_50_ / IC_50_ (or EC_50_).

### Anti-HCoV-229E and HCoV-OC43 compound testing

Human lung fibroblast MRC5 cells were cultured in a 96-well plate at 10,000 cells/well in EMEM+10% FBS overnight. The next day the culture medium was removed from each well and compounds in EMEM + 2% FBS + 0.5% DMSO were added. Tanespimycin, Bortezomib and Homoharringtonine were tested using 2 *μ*M starting concentration, 8-point, 3-fold serial dilution. After a 1-hour pre-incubation with testing compounds, HCoV-229E or HCoV-OC43 viruses at an MOI of 0.01 were added to the culture with compounds for another 1 hour. Cell-Titer Glo assay was added after additional incubation time with compounds of interest (HCoV-229E for 96 hours and HCoV-OC43 for 120 hours). Assay readout was performed using luminescence measurement on a Tecan Spark plate reader.

CC_50_ values were determined by applying nonlinear fit of luminescence readouts from uninfected, compound-treated wells against compound concentration. Likewise, EC_50_ values were determined by nonlinear fit of luminescence readouts from virus-infected, compound-treated wells. Data analysis was performed using GraphPad Prism 8, and SI calculated as the ratio of CC_50_ to EC_50_.

### Clinical dosing estimation of compounds

| Drug | Anti-Covid IC<sub>50</sub> | Known Pharmacology | Clinical Exposure | Reference |
| --- | --- | --- | --- | --- |
| Lacidipine<sup>1</sup> | 14.89 μM | Calcium channel blocker | 0.003 – 0.013 μM | [26] |
| Trametinib<sup>2</sup> | 14.95 μM | MEK1/2 inhibitor | 0.045 μM | [27] |
| Homoharringtonine<sup>3</sup> | 0.16 μM | Inhibition of protein synthesis | 0.046 μM | [28] |
| Bortezomib<sup>4</sup> | 1.39 μM | Reversible inhibitor of the chymotrypsin-like activity of the 26S proteasome | 0.29 μM | [29] |
| <sup>1</sup>Cmax after a single dose of 4mg per individual ranged from 1.6 to 5.7 ng/mL; molecular weight is 455.5 ng/nmol [26].<br><sup>2</sup>Cmax after a single dose of 2 mg per individual is 27.6 ng/mL; molecular weight is 615 ng/nmol [27].<br><sup>3</sup>Cmax after a single dose of 1.25 mg/m<sup>2</sup> is: 25.1 ng/mL; molecular weight is 545.6 ng/nmol [28].<br><sup>4</sup>Cmax after the first dose of 1.3 mg/m<sup>2</sup> is: 112 ng/mL; molecular weight is 384.24 ng/nmol [29]. |  |  |  |  |

## Competing Interests

SARS-CoV-2 studies were conducted at Institut Pasteur Korea (South Korea) and funded by Arum Therapeutics and Auransa Inc. HCoV-229E and HCoV-OC43 studies were conducted and funded by Experimental Drug Development Centre (Singapore). Auransa Inc. sourced all compounds used in *in vitro* testing.

Shen Z., Guo J., Song G., Louie B., Halberg A., Luedtke G.R., Protter A., Visuthikraisee V., and Lum P.Y. are employees of Auransa Inc. and hold equity in the company.

Baik T. is an employee of Arum therapeutics and holds equity in the company.

Fong J.Y. and Koh X. are employees of Experimental Drug Development Centre (Singapore) and have no conflict of interest.

## Author Contributions

The authors confirm contribution to the paper as follows:

<u>*In silico* data curation and analysis:</u> Shen Z., Guo J., Song G., Louie B., Visuthikraisee V., Lum P.Y.

<u>Compound selection:</u> Halberg A., Shen Z., Guo J., Luedtke G.R., Louie B., Protter A., Baik T., Lum P.Y.

<u>*In vitro* experiment and interpretation:</u> Halberg A., Fong J.Y., Koh X., IPK South Korea, Shen Z., Guo J., Lum P.Y.

<u>Compound clinical exposure estimation:</u> Protter A.

<u>Manuscript preparation:</u> Shen Z., Guo J., Halberg A., Lum P.Y.

## Notes

### Summary of Updates

The superscript footnotes for clinical exposure table's "Clinical Exposure" column have been removed. Header "Reference" has been changed to "References".

## References

1. Wang X, Zhou Y, Liu L, Ma J, Wu H, Zhao L, et al. Coronavirus disease 2019 (COVID-19): diagnosis and prognosis. Blood Genomics. 2020;4: 96–107. doi:10.46701/BG.2020022020120

2. Li X, Chang J, Chen S, Wang L, Yau TO, Zhao Q, et al. Genomic Feature Analysis of Betacoronavirus Provides Insights Into SARS and COVID-19 Pandemics. Front Microbiol. 2021;12. Available: https://www.frontiersin.org/article/10.3389/fmicb.2021.614494

3. Fight the Pandemic: Treatments. In: Artis Ventures [Internet]. [cited 7 Mar 2022]. Available: https://www.av.co/covid-treatments

4. Jeon S, Ko M, Lee J, Choi I, Byun SY, Park S, et al. Identification of Antiviral Drug Candidates against SARS-CoV-2 from FDA-Approved Drugs. Antimicrob Agents Chemother. 2020;64: e00819–20. doi:10.1128/AAC.00819-20

5. Choy K-T, Wong AY-L, Kaewpreedee P, Sia SF, Chen D, Hui KPY, et al. Remdesivir, lopinavir, emetine, and homoharringtonine inhibit SARS-CoV-2 replication in vitro. Antiviral Res. 2020;178: 104786. doi:10.1016/j.antiviral.2020.104786

6. Zhang H, Yang Y, Li J, Wang M, Saravanan KM, Wei J, et al. A novel virtual screening procedure identifies Pralatrexate as inhibitor of SARS-CoV-2 RdRp and it reduces viral replication in vitro. PLOS Comput Biol. 2020;16: e1008489. doi:10.1371/journal.pcbi.1008489

7. Gao G, Luo H. The ubiquitin–proteasome pathway in viral infectionsThis paper is one of a selection of papers published in this Special Issue, entitled Young Investigator’s Forum. Can J Physiol Pharmacol. 2006;84: 5–14. doi:10.1139/y05-144

8. Pan X, Li X, Ning S, Zhi H. Inferring SARS-CoV-2 functional genomics from viral transcriptome with identification of potential antiviral drugs and therapeutic targets. Cell Biosci. 2021;11: 171. doi:10.1186/s13578-021-00684-4

9. Adhami M, Sadeghi B, Rezapour A, Haghdoost AA, MotieGhader H. Repurposing novel therapeutic candidate drugs for coronavirus disease-19 based on protein-protein interaction network analysis. BMC Biotechnol. 2021;21: 22. doi:10.1186/s12896-021-00680-z

10. Xing J, Shankar R, Drelich A, Paithankar S, Chekalin E, Dexheimer T, et al. Analysis of Infected Host Gene Expression Reveals Repurposed Drug Candidates and Time-Dependent Host Response Dynamics for COVID-19. Genomics; 2020 Apr. doi:10.1101/2020.04.07.030734

11. Schubert K, Karousis ED, Jomaa A, Scaiola A, Echeverria B, Gurzeler L-A, et al. SARS-CoV-2 Nsp1 binds the ribosomal mRNA channel to inhibit translation. Nat Struct Mol Biol. 2020;27: 959–966. doi:10.1038/s41594-020-0511-8

12. Chen Y, Lear TB, Evankovich JW, Larsen MB, Lin B, Alfaras I, et al. A high-throughput screen for TMPRSS2 expression identifies FDA-approved compounds that can limit SARS-CoV-2 entry. Nat Commun. 2021;12: 3907. doi:10.1038/s41467-021-24156-y

13. Huang C-T, Chao T-L, Kao H-C, Pang Y-H, Lee W-H, Hsieh C-H, et al. Enhancement of the IFN-β-induced host signature informs repurposed drugs for COVID-19. Heliyon. 2020;6: e05646. doi:10.1016/j.heliyon.2020.e05646

14. Ma H, Wen H, Qin Y, Wu S, Zhang G, Wu C-I, et al. Homo-harringtonine, highly effective against coronaviruses, is safe in treating COVID-19 by nebulization. Sci China Life Sci. 2022 [cited 12 Apr 2022]. doi:10.1007/s11427-021-2093-2

15. Li C, Chu H, Liu X, Chiu MC, Zhao X, Wang D, et al. Human coronavirus dependency on host heat shock protein 90 reveals an antiviral target. Emerg Microbes Infect. 2020;9: 2663–2672. doi:10.1080/22221751.2020.1850183

16. Jang WD, Jeon S, Kim S, Lee SY. Drugs repurposed for COVID-19 by virtual screening of 6,218 drugs and cell-based assay. Proc Natl Acad Sci. 2021;118: e2024302118. doi:10.1073/pnas.2024302118

17. Beigel JH, Tomashek KM, Dodd LE, Mehta AK, Zingman BS, Kalil AC, et al. Remdesivir for the Treatment of Covid-19 — Final Report. N Engl J Med. 2020;383: 1813–1826. doi:10.1056/NEJMoa2007764

18. Eastman RT, Roth JS, Brimacombe KR, Simeonov A, Shen M, Patnaik S, et al. Remdesivir: A Review of Its Discovery and Development Leading to Emergency Use Authorization for Treatment of COVID-19. ACS Cent Sci. 2020;6: 672–683. doi:10.1021/acscentsci.0c00489

19. Warren TK, Jordan R, Lo MK, Ray AS, Mackman RL, Soloveva V, et al. Therapeutic efficacy of the small molecule GS-5734 against Ebola virus in rhesus monkeys. Nature. 2016;531: 381–385. doi:10.1038/nature17180

20. Tchesnokov EP, Feng JY, Porter DP, Götte M. Mechanism of Inhibition of Ebola Virus RNA-Dependent RNA Polymerase by Remdesivir. Viruses. 2019;11: E326. doi:10.3390/v11040326

21. Large clinical trial to study repurposed drugs to treat COVID-19 symptoms. In: National Institutes of Health (NIH) [Internet]. 19 Apr 2021 [cited 7 Mar 2022]. Available: https://www.nih.gov/news-events/news-releases/large-clinical-trial-study-repurposed-drugs-treat-covid-19-symptoms

22. Patro R, Duggal G, Love MI, Irizarry RA, Kingsford C. Salmon provides fast and bias-aware quantification of transcript expression. Nat Methods. 2017;14: 417–419. doi:10.1038/nmeth.4197

23. Subramanian A, Tamayo P, Mootha VK, Mukherjee S, Ebert BL, Gillette MA, et al. Gene set enrichment analysis: A knowledge-based approach for interpreting genome-wide expression profiles. Proc Natl Acad Sci. 2005;102: 15545–15550. doi:10.1073/pnas.0506580102

24. Szklarczyk D, Gable AL, Nastou KC, Lyon D, Kirsch R, Pyysalo S, et al. The STRING database in 2021: customizable protein–protein networks, and functional characterization of user-uploaded gene/measurement sets. Nucleic Acids Res. 2021;49: D605–D612. doi:10.1093/nar/gkaa1074

25. Szklarczyk D, Gable AL, Lyon D, Junge A, Wyder S, Huerta-Cepas J, et al. STRING v11: protein–protein association networks with increased coverage, supporting functional discovery in genome-wide experimental datasets. Nucleic Acids Res. 2019;47: D607–D613. doi:10.1093/nar/gky1131

26. Lee CR, Bryson HM. Lacidipine. A review of its pharmacodynamic and pharmacokinetic properties and therapeutic potential in the treatment of hypertension. Drugs. 1994;48: 274–296. doi:10.2165/00003495-199448020-00010

27. Voon PJ, Chen EX, Chen HX, Lockhart AC, Sahebjam S, Kelly K, et al. Phase I pharmacokinetic study of single agent trametinib in patients with advanced cancer and hepatic dysfunction. J Exp Clin Cancer Res CR. 2022;41: 51. doi:10.1186/s13046-021-02236-7

28. Nemunaitis J, Mita A, Stephenson J, Mita MM, Sarantopoulos J, Padmanabhan-Iyer S, et al. Pharmacokinetic study of omacetaxine mepesuccinate administered subcutaneously to patients with advanced solid and hematologic tumors. Cancer Chemother Pharmacol. 2013;71: 35–41. doi:10.1007/s00280-012-1963-2

29. VELCADE_PRESCRIBING_INFORMATION.pdf. Available: https://www.velcade.com/files/pdfs/VELCADE_PRESCRIBING_INFORMATION.pdf

